# Development of additional microsatellite primers for the mangrove tree species *Avicennia germinans*

**DOI:** 10.1101/2020.12.14.422622

**Authors:** Hayley Craig, Ilka C. Feller, Jennifer K. Rowntree

## Abstract

**Objective:** The objective of this study was to develop additional microsatellite primers for the mangrove tree species *Avicennia germinans* that work in multiplex PCR panels to enable cost effective population analyses of this species at a finer scale.

**Results:** Primer sets were identified from whole genome sequencing data and combined into multiplex PCR panels. Five multiplex reactions containing 20 novel primer sets with trinucleotide repeats were successfully developed. Fifteen of the microsatellite loci were polymorphic in the samples tested, with 1-4 alleles per locus.

## Introduction

North-eastern Florida is currently considered the northern most range limit of mangrove distribution (Cavanaugh et al. 2019) and its mangrove ecosystems are poor in plant species diversity, with only three species found in the state (Spalding 2010). This makes these environments ideal for investigating a range of topics including: range expansion (Kennedy et al. 2017), effects of intra-specific diversity on ecosystem properties (Craig et al. 2020), and frost adaptation (Hayes et al. 2020). Of the three true mangrove tree species found in Florida, *Avicennia germinans* is the most wide-ranging (Nelson 2010) and several studies have used microsatellite markers to investigate population structure of this species at the regional scale (e.g. Hodel et al. 2016; Kennedy et al. 2020). While microsatellite markers have been developed for A. germinans (Cerón-Souza et al. 2006; Nettel et al. 2005; Mori et al. 2010), they have been designed for use in singleplex PCR reactions of varying cycle conditions. The cost and time constraints of running multiple PCRs have limited the number of markers used in studies to date. To enable researchers to look at finer scale population genetic structure of A. germinans, both in Florida and further afield, we have developed novel microsatellite markers using material collected from this region, and combined them into multiplex PCR panels to reduce time and costs needed for analysis. These new markers could be combined with existing markers to boost the total number used in studies and increase the resolution at which inferences can be drawn.

## Methods

### Microsatellite development

Microsatellite primers were developed following the Griffiths et al. (2016) workflow. Briefly, genomic DNA from one A. germinans individual collected from Pine Island was selected for sequencing using the MiSeq System (Illumina Inc., San Diego, USA) with 2 x 250 bp read length to create a paired-end library. Raw data (2 209 638 reads) were imported into Palfinder Galaxy, a customised version of Galaxy (Afgan et al. 2016) managed by The University of Manchester’s Bioinformatics Core Facility. Reads were filtered and trimmed based on quality using Trimmomatic (Bolger et al. 2014), microsatellites were isolated using Pal_finder (Castoe et al. 2012), primers were designed using Primer3 (Koressaar & Remm 2007; Untergasser et al. 2012), and primers were then filtered using Pal_filter (Griffiths et al. 2016) and PANDAseq (Masella et al. 2012). Primer design was optimised to work with the Type-it Microsatellite PCR kit (QIAGEN, Hilden, Germany).

Pal_filter identified primers for 143 loci in regions containing microsatellites with tri- or tetranucleotide perfect repeat motifs. We excluded dinucleotide repeats because although they can detect greater genetic variation (Merritt et al. 2015), they are more prone to slipped-strand mispairing (stuttering), which can result in genotyping errors (Zalapa et al. 2012). As only 20 primer sets were identified in the final assembly step of the pipeline with PANDAseq, an additional 10 primers were randomly selected from the filtered primers (without assembly) output for testing. Primers were tested following the methods developed by Culley et al. (2013), adjusted for 5 μl reactions instead of 10 μl. In summary, unlabelled and untailed primer pairs were first tested in singleplex PCRs using the Type-it Microsatellite PCR kit to check whether they amplified fragments. All primers were tested in the same conditions to find those that could be combined together into multiplexes. All PCR amplifications were carried out under the following conditions: 95 °C for 5 min; followed by 28 cycles of 95 °C for 30 s, 60 °C for 1 min 30 s and 72 °C for 30 s; followed by 30 min extension at 60 °C. PCR product was examined by 1.5% (w/v) agarose 0.5x TBE buffer (45mM Tris-borate, 1mM EDTA) gel electrophoresis with GelGreen stain (Biotium Inc., Fremont, CA, USA).

Twenty three of the 30 primer sets tested amplified successfully and continued to the second stage of testing; amplifying forward tailed (but unlabelled) primers with reverse primers in singleplex PCR. Twenty of the primer sets amplified successfully and continued to the third stage of testing in singleplex PCRs with a fluorescently labelled tail added. 0.5 μl of labelled PCR product was mixed with 9.1 μl HiDi formamide (Thermo Fisher Scientific, Waltham, MA, USA) and 0.4 μl LIZ600 sizing ladder (Applied Biosystems, Foster City, CA, USA) prior to fragment sizing on the ABI3730 (Applied Biosystems, Foster City, CA, USA).

Once the size of the fragments from each locus was known, Multiplex Manager (Holleley & Geerts 2009) was used to combine loci into suitable PCR multiplexes using three fluorescent labels (6-FAM, HEX and PET). If any primer sets did not successfully amplify in multiplex for microsatellite scoring, multiplex combinations were optimised until all included primer pairs were working. Five multiplex PCR panels comprising 20 primer sets were tested (Table 1). All primer sets in final multiplexes were for trinucleotide repeat motifs.

**Table 1.**
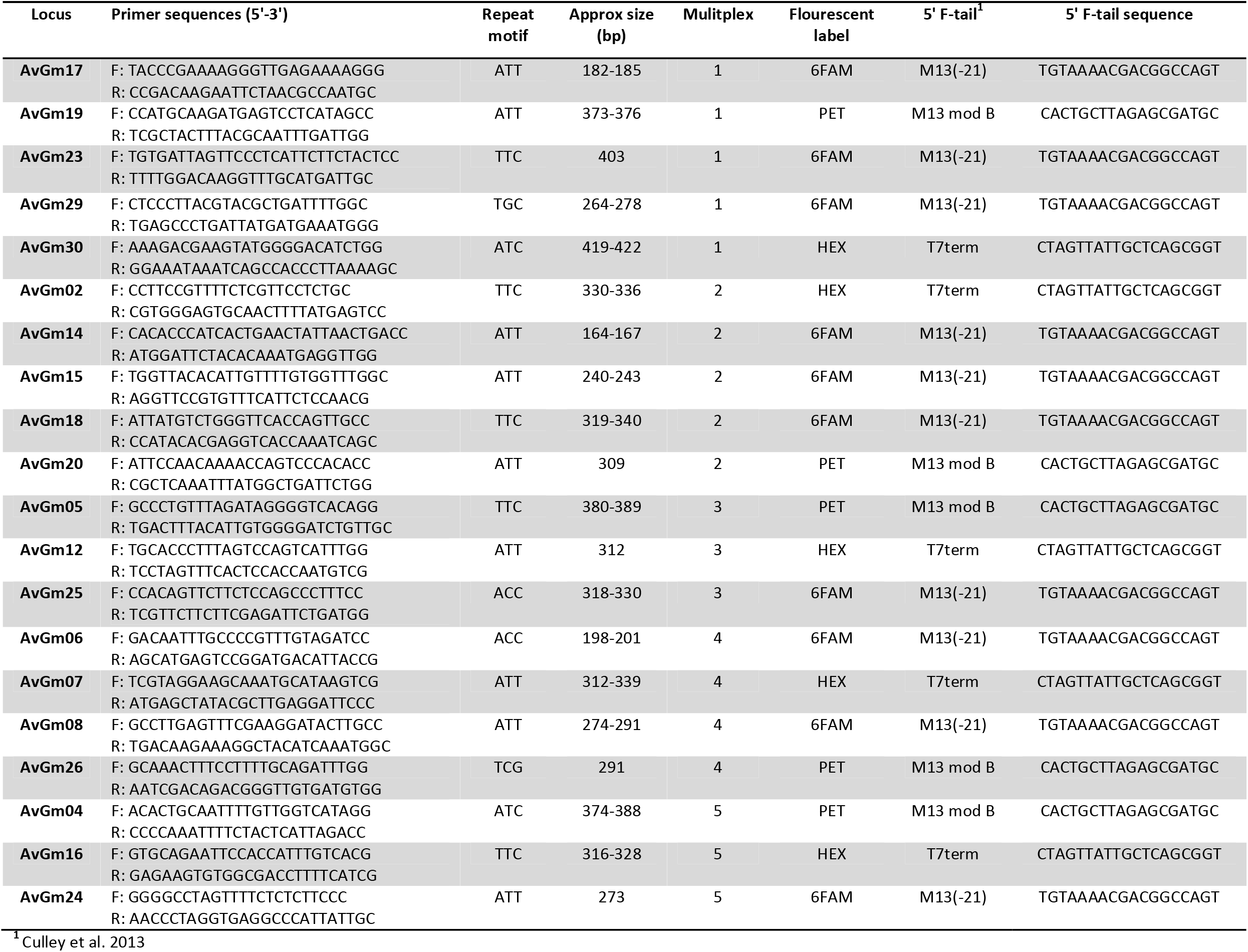
Characteristics of five microsatellite multiplex panels including 20 novel primer sets developed for *Avicennia germinans*

### Multiplex testing

Initial microsatellite primer testing was conducted on DNA extracted from the leaves of 16 A. germinans individuals from two sites within the Indian River Lagoon system: eight collected from Pine Island Conservation Area, Brevard County, Florida (28.4827 °N, 80.7240 °W) and eight from Avalon State Recreation Area on North Hutchinson Island, St Lucie County, Florida (27.5488 °N, 80.3269 °W). Genomic DNA was extracted from 30 mg dried, ground leaf tissue using the Qiagen DNeasy Plant extraction kit as per the manufacturer’s protocol. Quality and quantity of DNA extractions were checked using a NanoDrop 1000 spectrophotometer (Thermo Fisher Scientific, Wilmington, DE, USA). Once the multiplex PCR panels of microsatellites had been successfully optimised, final testing was performed on additional samples totalling 30 individuals from each of the two sites.

The microsatellite data were analysed using GeneMapper version 4.1 (Applied Biosystems, Foster City, CA, USA) to separate fragments based on dye colour and size and then to pick peaks for each locus. Some manual editing was required of bins due to differences observed in duplicate samples between plates. Binned data was imported into GenAlEx 6.502 (Peakall & Smouse 2012), to assess numbers of alleles, heterozygosity, and Hardy-Weinberg equilibrium.

## Results and Discussion

Overall, 15 of the 20 primer sets in the multiplex panels were found to be polymorphic within the samples from primer development tests (Tables 1 and 2). We identified 43 total alleles, number of alleles per locus ranged from 1–4, and 13 private alleles were identified. All loci conformed to Hardy–Weinberg equilibrium (HWE) at the Avalon site, but six loci deviated from Hardy-Weinberg equilibrium in the Pine Island samples.

**Table 2.** Genetic diversity of 20 microsatellites developed for *Avicennia germinans*

| Locus | Pine Island, FL (n = 30) |  |  | Avalon, FL (n = 30) |  |  |
| --- | --- | --- | --- | --- | --- | --- |
|  | A | H<sub>o</sub> | H<sub>e</sub> | A | H<sub>o</sub> | H<sub>e</sub> |
| AvGm02 | 2 | 0.033 | 0.033 | 3 | 0.367 | 0.455 |
| AvGm04 | 2 | 0.034 | 0.034 | 2 | 0.067 | 0.064 |
| AvGm05 | 3 | 0.233 | 0.383 | 3 | 0.500 | 0.546 |
| AvGm06 | 2 | 0.000* | 0.069 | 2 | 0.267 | 0.320 |
| AvGm07 | 2 | 0.179* | 0.484 | 3 | 0.367 | 0.415 |
| AvGm08 | 2 | 0.107 | 0.101 | 2 | 0.133 | 0.180 |
| AvGm12 | 1 | 0.000 | 0.000 | 1 | 0.000 | 0.000 |
| AvGm14 | 1 | 0.000 | 0.000 | 2 | 0.233 | 0.206 |
| AvGm15 | 2 | 0.067 | 0.064 | 1 | 0.000 | 0.000 |
| AvGm16 | 2 | 0.034* | 0.098 | 3 | 0.333 | 0.315 |
| AvGm17 | 1 | 0.000 | 0.000 | 2 | 0.067 | 0.064 |
| AvGm18 | 2 | 0.133* | 0.278 | 4 | 0.667 | 0.564 |
| AvGm19 | 2 | 0.167* | 0.299 | 2 | 0.467 | 0.500 |
| AvGm20 | 1 | 0.000 | 0.000 | 1 | 0.000 | 0.000 |
| AvGm23 | 1 | 0.000 | 0.000 | 1 | 0.000 | 0.000 |
| AvGm24 | 1 | 0.000 | 0.000 | 1 | 0.000 | 0.000 |
| AvGm25 | 1 | 0.000 | 0.000 | 2 | 0.033 | 0.033 |
| AvGm26 | 1 | 0.000 | 0.000 | 1 | 0.000 | 0.000 |
| AvGm29 | 2 | 0.033* | 0.095 | 3 | 0.467 | 0.413 |
| AvGm30 | 1 | 0.000 | 0.000 | 2 | 0.033 | 0.033 |

Note: A = number of alleles; H<sub>e</sub> = expected heterozygosity; H<sub>o</sub> = observed heterozygosity; n=number of individuals sampled
\* Significant deviation from Hardy–Weinberg equilibrium (p < 0.05)

Here, we have successfully developed novel microsatellite markers for the mangrove tree species A. germinans that work in multiplex PCR conditions. These could be combined with existing markers from the literature for more in-depth population analyses in areas of potentially low genetic diversity.

## Authors’ contributions

HC and JKR conceived and designed the study. HC collected the field samples, designed and tested the microsatellite primers, analysed the data, and prepared the manuscript. JKR and ICF supervised the project and provided guidance on the analyses. All authors read and approved the final manuscript.

## Acknowledgements

The study contained in this publication was supported by Natural Environment Research Council (NERC) EAO Doctoral Training Partnership (NE/L002469/1) funding to HC. The authors wish to thank John Paul Kennedy for his assistance in the field and useful insights. Microsatellite fragment sizing was conducted at The University of Manchester Genomic Technologies Core Facility by Paul Fullwood and Fraser Combe. The authors would also like to thank Florida State Parks and the Brevard County Parks and Recreation Department Environmentally Endangered Lands Program for access to the sites and permission to collect samples.

## Notes

### Competing Interest Statement

The authors have declared no competing interest.

